# Deconvolution reveals compositional differences in high-grade serous ovarian cancer subtypes

**DOI:** 10.1101/2023.06.14.544991

**Authors:** Ariel A. Hippen, Natalie R. Davidson, Mollie E. Barnard, Lukas M. Weber, Jason Gertz, Jennifer A. Doherty, Stephanie C. Hicks, Casey S. Greene

## Abstract

Ovarian cancer is a deadly disease with few effective therapies. The most common form is high-grade serous ovarian cancer (HGSOC). Transcriptomic subtypes of HGSOC have shown promise in characterizing tumor heterogeneity and are associated with survival. Gene expression signatures for the subtypes suggest variation in stromal cell types in the tumor microenvironment (TME). Here, we characterize the TME composition of HGSOC on a population scale by performing deconvolution on bulk transcriptomic data. We use comprehensive cell type profiles from 164 HGSOC tumor samples from two independent reference datasets, in order to compare cell type proportions across and within bulk transcriptomic datasets, and assess their alignment to the subtypes proposed by The Cancer Genome Atlas. We also assess the relationship between tumor composition and clinical outcomes. Our results suggest that HGSOC transcriptomic subtypes are driven by TME composition, specifically fibroblast and immune cell content, and we propose a modified HGSOC subtype model informed by cell composition.

## Introduction

Ovarian cancer is the fifth-most common cause of cancer death for women in the US, responsible for almost 13,000 deaths annually [1]. High-grade serous ovarian cancer (HGSOC) is the most common form, accounting for 70% of cases and the majority of deaths [2]. Most patients initially respond well to standard-of-care chemotherapy, but many tumors quickly recur and become treatment-resistant. Few new therapies have been introduced in the past decade [3]. While some genetic contributions to clinical response have been well-characterized, like the success of PARP inhibitors in BRCA-mutated tumors [4], much of the heterogeneity in treatment outcomes is still being characterized.

Transcriptomic subtypes are a useful model of tumor heterogeneity in some cancers. For instance, the PAM50 breast cancer subtypes have prognostic value independent of other clinical and histological factors [5, 6]. Contributors to The Cancer Genome Atlas (TCGA) identified four transcriptomic subtypes of HGSOC: mesenchymal, differentiated, immunoreactive, and proliferative [7]. While the most appropriate number of transcriptomic subtypes is disputed [8, 9], studies of subtypes across many populations have demonstrated differential survival [10–13], suggesting that subtype features reflect real intertumoral differences.

A noteworthy difference among transcriptomic subtypes of HGSOC is the variation in the prevalence of stromal cell types in the tumor microenvironment (TME). One of the earliest papers describing molecular subtypes in HGSOC found that a high stromal gene signature correlated with worse survival outcomes [10]. This trend has been replicated and associated with the mesenchymal subtype as described by TCGA [11, 14, 15]. The immunoreactive subtype, so named for the high expression of immune-associated genes, is associated with better survival [16]. More recent papers have suggested that molecular subtype classification is not merely associated with cell composition but rather is driven by it. In a simulation study, increasing the tumor stromal admixture was enough to change the subtype assignment of a bulk tumor [17]. When performing subtype classification on single cells, cell assignments were largely grouped by cell type [18], and subtype gene signatures were highly correlated with cell types [19]. These results suggest that the variable cellular composition of bulk tumors could be a confounding factor in the assessment of HGSOC heterogeneity [20].

Here, we apply a holistic approach to characterizing the composition of the tumor microenvironment of HGSOC on hundreds of samples, allowing us to make population-level inferences. We do this by performing deconvolution of several HGSOC datasets profiled using bulk transcriptomics. Broadly, deconvolution describes breaking up a matrix of data into proportions of discrete component parts—in this case, gene expression count data into proportions of component cell types in a bulk transcriptomic sample [21]. A variety of deconvolution methods exist, with many of the latest using labeled single-cell RNA-seq (scRNA-seq) data as a reference profile exemplifying the gene expression of relevant cell types [22–24]. Here we use one such method, BayesPrism, to perform the deconvolution of HGSOC data [25]. We found in a previous study [26] that this method performed accurately on simulated data, and returned consistent results on real data with varying technical factors. BayesPrism is designed for use on cancer data, controlling for the high inter-sample heterogeneity observed in cancer cell gene expression [27, 28].

Using deconvolution, we obtain a comprehensive cell type profile for hundreds of bulk HGSOC samples, with estimates of proportions of all the cell types observed in scRNA-seq data from the same tumor type. We compare the cell type proportions across and within datasets, and stratify tumors based on their alignment to one of the TCGA transcriptomic subtype signatures [8]. We also assess the association between tumor composition and clinical outcomes, specifically success of surgical resection and overall survival. Finally, we propose a modified framework informed by cell composition for using HGSOC subtypes.

## Results

### HGSOC deconvolution results are concordant across datasets, reference profiles, and platforms

We deconvolved three datasets of bulk-profiled HGSOC tumors: those sequenced by TCGA using RNA-seq, those sequenced by TCGA using microarray profiling, and the samples from Tothill et al. [10] (Table 1). We performed analyses in parallel using different reference profiles: HGSOC-Penn/Utah data [29, 30], and data sampled from Vázquez-García et al [31]. Each reference profile comprised gene expression signatures from scRNA-seq data annotated by cell type (see Methods).

**Table 1.** Bulk datasets used as inputs for deconvolution.

| Name | Data type | Samples | Genes used |
| --- | --- | --- | --- |
| TCGA RNA-seq | RNA-seq | 300 | 15490 |
| TCGA Microarray | Microarray | 480 | 11074 |
| Tothill | Microarray | 147 | 15548 |

We first performed bulk deconvolution using the HGSOC-Penn/Utah single-cell reference profile (Fig. 1A-C). We observed certain consistent trends across all three bulk transcriptomic datasets; for instance, the composition of all tumors is predominated by epithelial cells, consistent with an epithelial cancer [32]. In all datasets, the second most common cell type is fibroblasts, with high inter-sample variability in fibroblast content. Samples in each dataset also had a smaller but non-trivial fraction of endothelial cells and macrophages.

**Fig 1.**
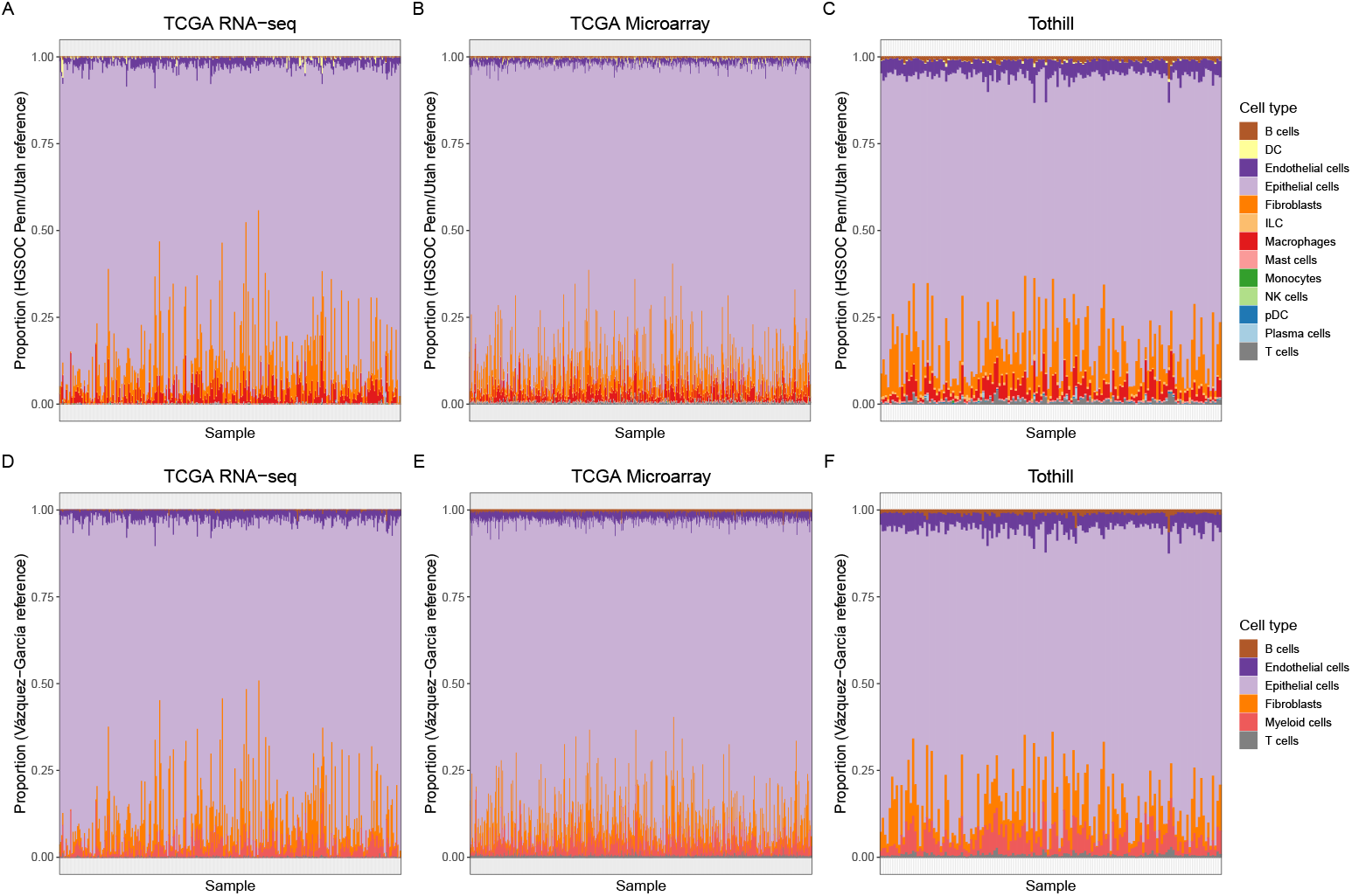
Deconvolution estimates. **(A-C)** Cell type proportion estimates output from BayesPrism when using the HGSOC-Penn/Utah reference profile. **(D-F)** Proportion estimates when using the Vázquez-García reference profile.

While there are prevailing similarities in the composition of tumors across datasets, some differences emerge. Both TCGA datasets, RNA-seq and microarray, had higher average epithelial content compared to Tothill (86.9% for TCGA microarray and 85.7% for TCGA RNA-seq compared to 79.1% for Tothill). This likely derives from a difference in sample selection across the datasets; TCGA only included samples with greater than 70% estimated tumor purity [7], whereas Tothill included samples with as low as 30% estimated tumor purity [10]. In turn, Tothill had a higher average content of most other cell types, particularly endothelial cells.

To assess whether deconvolution results were affected by the reference profile used, we performed parallel analyses using a representative 10% sample of single-cell data from Vázquez-García et al [31]. This reference profile was grouped into fewer cell types by the original authors than the HGSOC-Penn/Utah reference profile, meaning an exact comparison was not possible, but the deconvolution estimates were qualitatively similar in broad profiles (Fig. 1D-F). Matching cell types were highly correlated across samples; the Spearman rank-order correlation of fibroblast estimates across samples was greater than 0.986 for all datasets, and the Spearman correlation of epithelial cells was greater than 0.989 (Fig. 2A-C). From the qualitative and quantitative similarities, we infer that the tumor composition estimates do not appear to be dependent on the single-cell reference profile used.

**Fig 2.**
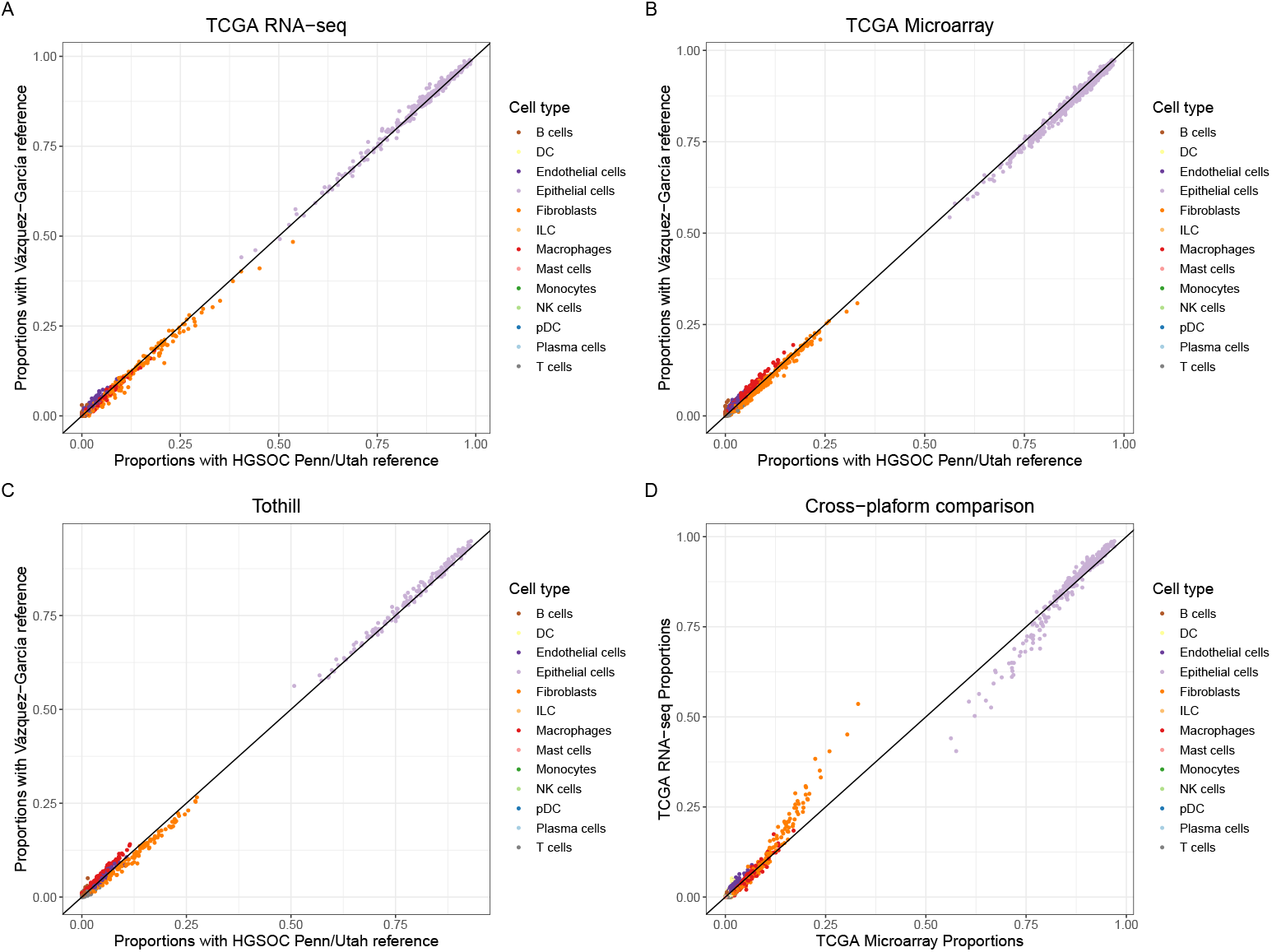
Correlation of cell type proportion estimates across reference profile and platform type. **(A-C)** The correlation of deconvolution output results between the HGSOC-Penn/Utah reference profile and the Vázquez-García reference profile for matching cell types. For completeness, here the estimate for “myeloid cells” in Vázquez-García is compared to the estimate for macrophages, the most abundant myeloid cell type, in HGSOC-Penn/Utah. **(D)** Cell type proportion estimates for TCGA samples sequenced using both RNA-seq and microarray (n = 255).

BayesPrism was designed for RNA-seq data [25] and to our knowledge has not previously been run on microarray datasets. To assess the validity of BayesPrism deconvolution of datasets profiled using microarrays, we compared the cell composition estimates for the 255 TCGA samples for which both microarray and RNA-seq data were available. The Pearson correlation for cell type proportions in each sample was very high, with an average correlation value of *r*=0.998 and all samples having a correlation value greater than *r*=0.905 (Fig. 2D). The biggest difference observed in the samples with relatively low correlation values was that the RNA-seq samples tended to estimate higher fibroblast content and consequently lower epithelial content than the same samples’ microarray estimates. Whether this is indicative of a global difference in microarray profiling vs. RNA-seq profiling or an artifact of differences in the TCGA workflow specifically remains to be seen. However, when comparing the estimates across samples in the dataset, the rank-ordering of samples was concordant for highly expressed cell types; the Spearman rank-order correlation of epithelial content and fibroblast content across samples was 0.984 and 0.993 respectively. This suggests that BayesPrism is a feasible way to estimate cell type composition of samples that have been characterized on a one-color microarray, particularly for within-dataset relative comparisons about the effect of cell type fraction on other biological or clinical factors such as survival.

### Variations in tumor composition support a model of three transcriptional subtypes

As stated previously, there is a wealth of literature suggesting a relationship between tumor composition and the four transcriptional HGSOC subtypes originally reported by TCGA. We used the pipeline introduced in [8] to assign samples in each dataset to transcriptional subtypes via K-means clustering (see Methods). We then assessed the relationship between cell proportion estimates and the predicted transcriptional subtype across bulk datasets.

In each dataset, samples assigned to the mesenchymal subtype had a substantially greater percentage of fibroblasts than the other subtypes (Fig. 3A). This aligns with previous work linking the mesenchymal subtype gene signature to fibroblast gene expression [19, 33]. Across all datasets, the mesenchymal subtype had the lowest percentage of malignant epithelial cells (commensurate with its increased fibroblast content) and the differentiated subtype the highest (Fig. 3B). While the total proportion of endothelial cells varied across datasets, we did not observe a difference in endothelial content between subtypes (Fig. 3C).

**Fig 3.**
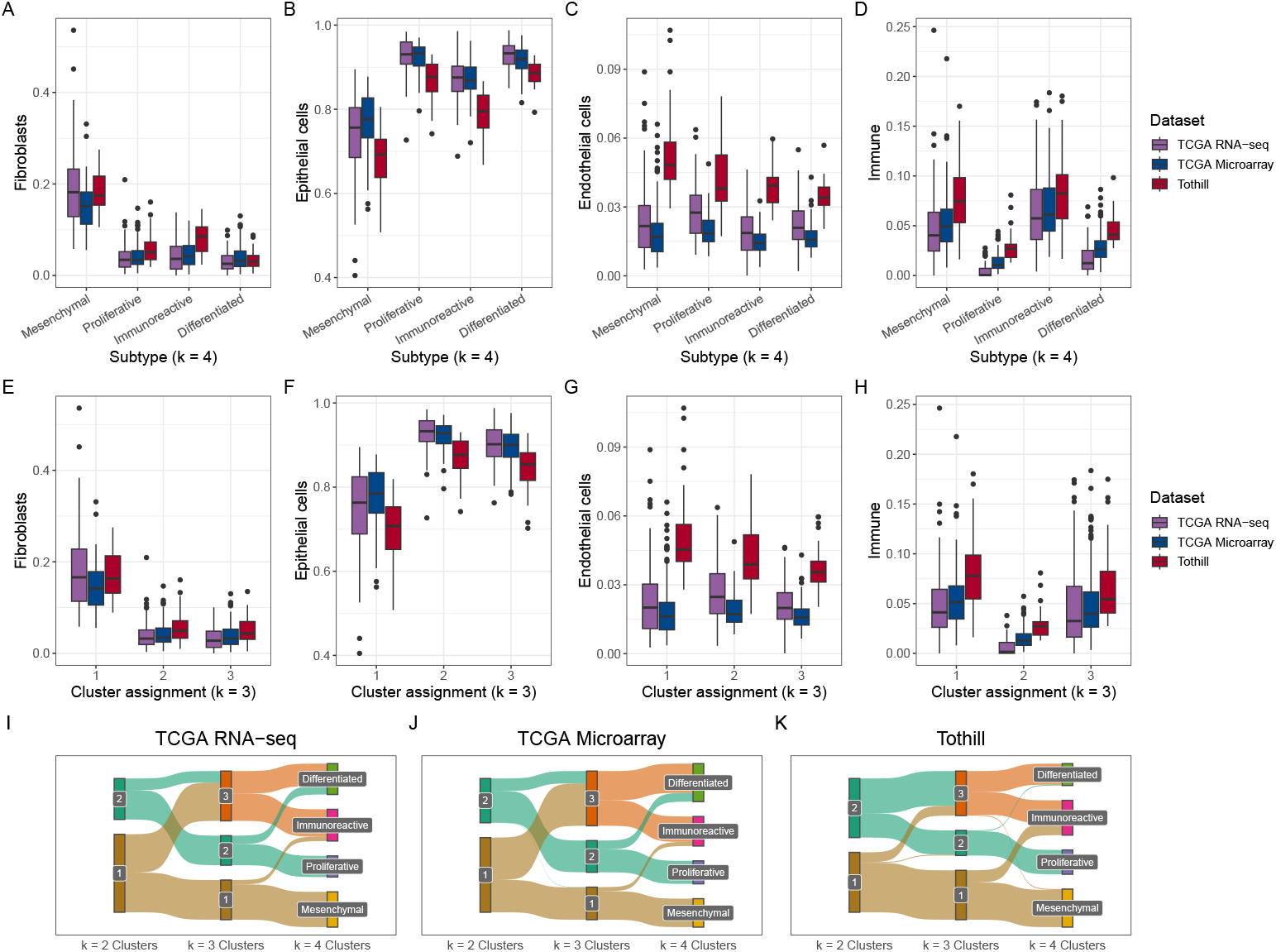
Cell type proportions across HGSOC transcriptional subtypes. We performed k-means clustering on each of the bulk transcriptomic datasets for *k* = 2, 3, and 4 clusters. **(A-D)** Cell type proportions stratified by cluster assignment at *k* = 4, mapped to the nomenclature of the original TCGA subtypes [7]. **(E-H)** Cell type proportions stratified by cluster assignment at *k* = 3. **(I-K)** Sankey diagrams showing how sample clustering differs across values of *k* in each datset.

Because we had many types of immune cells present in relatively small proportions in our bulk tumors, rather than comparing each cell type across subtypes individually, we combined all immune cell types (see Table 2) into a single immune score and compared that across subtypes. We observed that across all datasets studied, the mesenchymal and immunoreactive subtypes had a larger immune fraction than either the differentiated or proliferative subtype. Across all datasets, the proliferative subtype had the lowest immune composition (Fig. 3D).

**Table 2.**
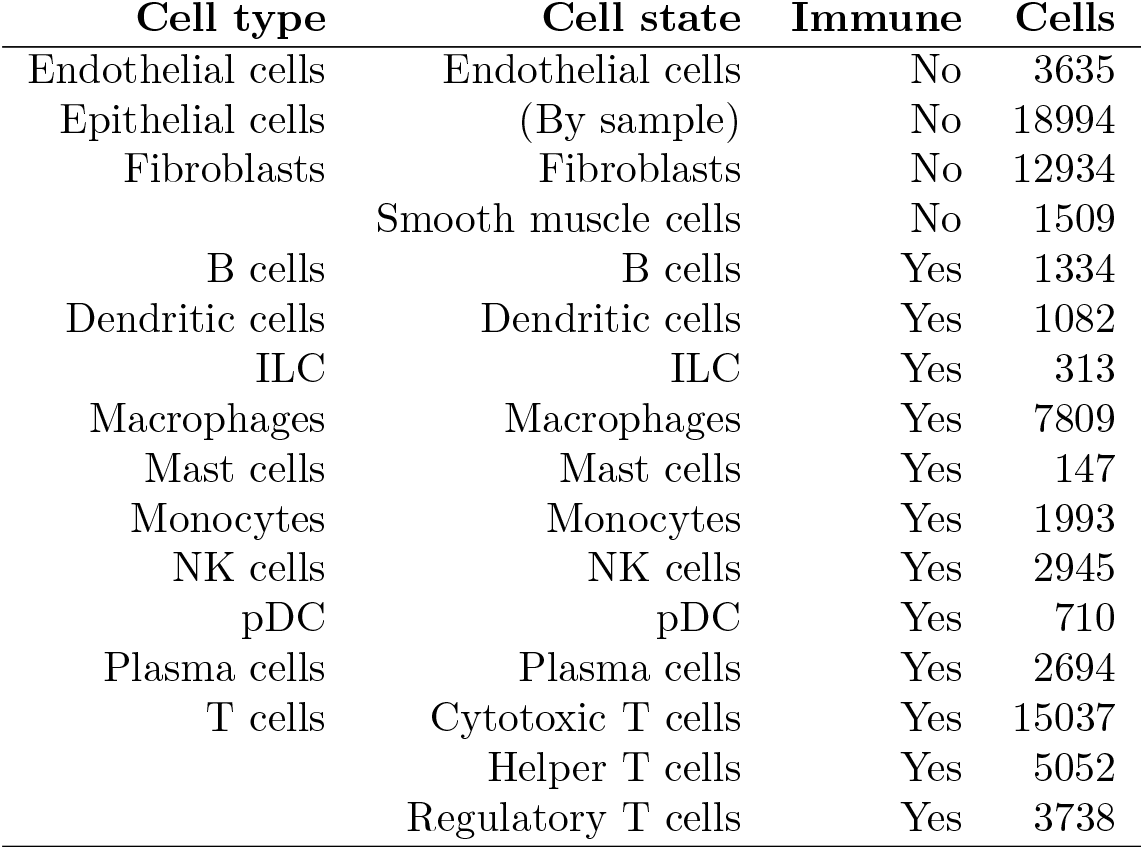
HGSOC-Penn/Utah single-cell data stratified by cell type and state.

| Cell type | Cell state | Immune | Cells |
| --- | --- | --- | --- |
| Endothelial cells | Endothelial cells | No | 3635 |
| Epithelial cells | (By sample) | No | 18994 |
| Fibroblasts | Fibroblasts | No | 12934 |
|  | Smooth muscle cells | No | 1509 |
| B cells | B cells | Yes | 1334 |
| Dendritic cells | Dendritic cells | Yes | 1082 |
| ILC | ILC | Yes | 313 |
| Macrophages | Macrophages | Yes | 7809 |
| Mast cells | Mast cells | Yes | 147 |
| Monocytes | Monocytes | Yes | 1993 |
| NK cells | NK cells | Yes | 2945 |
| pDC | pDC | Yes | 710 |
| Plasma cells | Plasma cells | Yes | 2694 |
| T cells | Cytotoxic T cells | Yes | 15037 |
|  | Helper T cells | Yes | 5052 |
|  | Regulatory T cells | Yes | 3738 |

The appropriate number of HGSOC transcriptional subtypes is disputed in the literature [8, 9, 13]. Because of this, we also used k-means clustering with *k* = 3 to assign all bulk samples to one of three subtypes, and compare cell composition across these subtypes (Fig. 3E-H). Again we found one subtype (subtype 1) with substantially higher fibroblast expression. This subtype also had relatively high immune content.

Another subtype (subtype 3) had high immune content but low fibroblast expression. A third subtype (subtype 2) had high epithelial expression, with neither high fibroblast nor immune content. Samples membership in clusters at *k* = 3 and *k* = 4 was largely consistent (and less so at *k* = 2). Across all datasets, samples in subtype 1 at *k* = 3 were mostly assigned to the mesenchymal subtype at *k* = 4, whereas samples in subtype 2 at *k* = 3 were mostly assigned to the proliferative subtype at *k* = 4. Samples in subtype 3 at *k* = 3 were from both the differentiated and the immunoreactive subtypes at *k* = 4 (Fig. 3I-K).

### Tumor composition explains some variation in clinical outcomes

Much of the interest in HGSOC transcriptional subtypes is related to their observed differences in length of survival [20]. Given the connection between transcriptional subtype and tumor composition we had already observed, we next assessed if tumor composition is associated with survival and other clinical outcomes. The most distinct difference we observed in tumor composition across samples and across subtypes was in the proportion of fibroblasts. Since the mesenchymal subtype has the highest proportion of fibroblasts and is also associated with the worst survival [11], we hypothesized that patients with a high fibroblast content in their samples would have a comparatively worse prognosis than patients with low fibroblast content. Fibroblast content was not normally distributed within datasets, with most samples in each dataset having a low or very low proportion of fibroblasts. We therefore defined samples as “high fibroblast” if they were in the top quartile of fibroblast content for their dataset (11.8% for TCGA RNA-seq, 9.9% for TCGA Microarray, 15.1% for Tothill) (Figure S1). We compared fibroblast status to survival by plotting Kaplan-Meier curves (Fig. 4A-D). In the TCGA RNA-seq and Tothill datasets, high fibroblast status was associated with a modestly worse prognosis, but we did not observe a similar association in the TCGA Microarray data. When considering all three datasets together, high fibroblast samples had lower survival than other samples at almost all time points (Fig. 4D).

**Fig 4.**
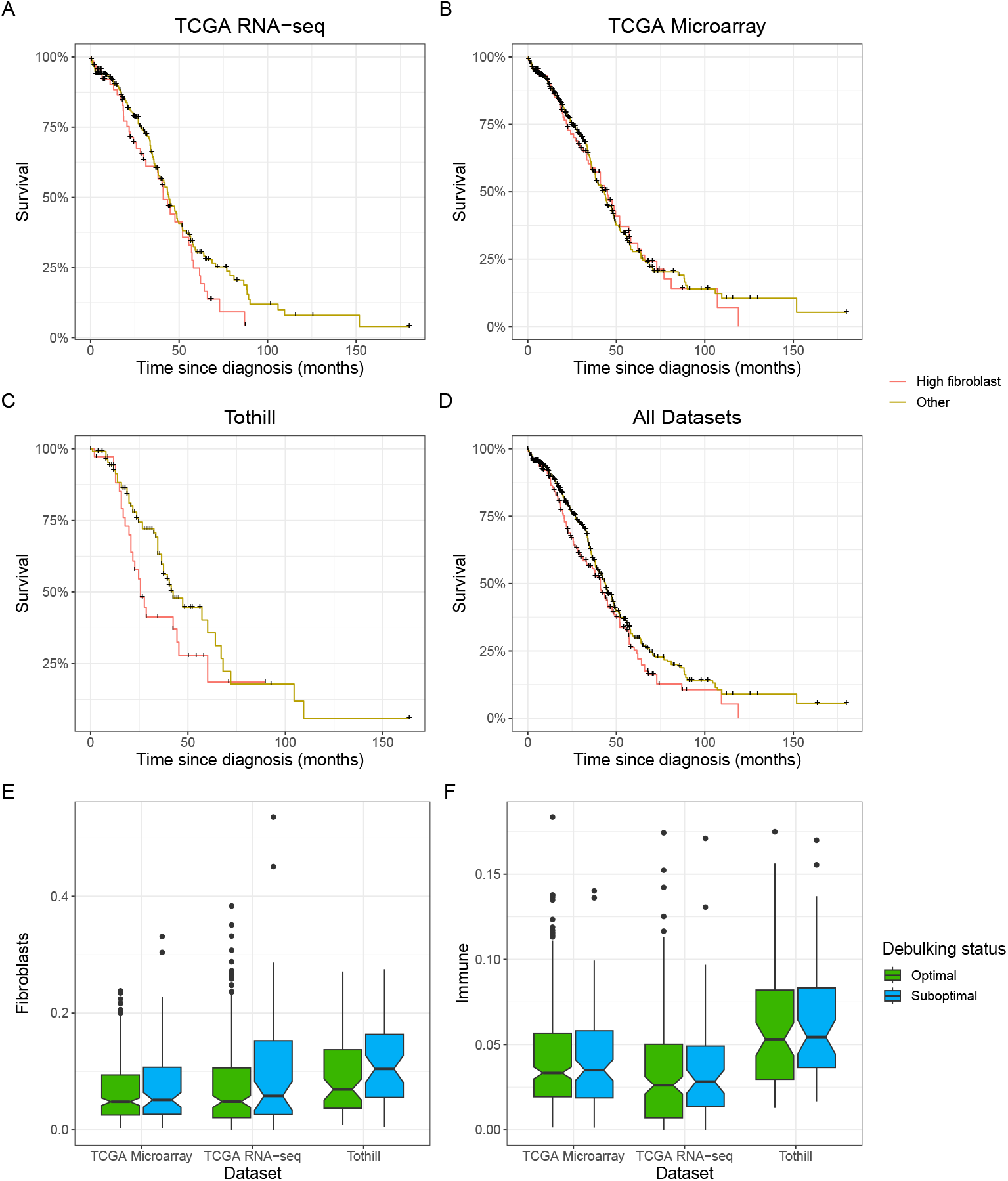
Tumor composition and clinical outcomes. **(A-D)** Kaplan-Meier survival curves stratified between high fibroblast tumors (top quartile of fibroblast content per dataset) and non-high fibroblast (all other quartiles). **(E)** Boxplot showing fraction of fibroblast content in a sample, stratified by dataset and debulking status (optimal vs. suboptimal). **(F)** Samples’ immune content stratified by dataset and debulking status.

We used Cox Proportional Hazards models to assess the risk of death based on high fibroblast status across all datasets. We ran the models two ways, once using only fibroblast content and once using covariates associated with HGSOC survival, namely age, tumor stage, and outcome of surgical debulking. When considered in univariate models, all datasets showed an increased risk of death for high fibroblast samples compared to non-high fibroblast samples (TCGA RNA-seq Hazard Ratio (HR)=1.36, 95% CI: 0.94-1.95; TCGA Microarray HR=1.05, 95% CI: 0.77-1.42; Tothill HR=1.57, 95% CI: 0.96-2.59). After controlling for age, tumor stage, and outcome of surgical debulking, there was essentially no association in TCGA (TCGA RNA-seq Hazard Ratio (HR)=1.04, 95% CI: 0.70-1.55; TCGA Microarray HR=0.97, 95% CI: 0.69-1.36), but the magnitude of the association increased in Tothill (HR=1.72, 95% CI: 0.98-3.01). The next greatest source of variability across samples after fibroblasts was variation in immune cell content. We performed the same analyses as with fibroblasts, comparing survival rates of samples in the top quartile of immune fraction vs. all others (threshold of 5.0% for TCGA RNA-seq. 5.7% for TCGA Microarray, 8.9% for Tothill) (Figure S1). We did not see consistent associations for high immune content on overall survival when considered in a univariate model (TCGA RNA-seq HR=1.21, 95% CI: 0.81-1.79; TCGA Microarray HR=1.00, 95% CI: 0.73-1.37; Tothill HR=0.88, 95% CI: 0.51-1.52) or after controlling for age, tumor stage, and outcome of surgical debulking (TCGA RNA-seq HR=1.47, 95% CI: 0.96-2.26; TCGA Microarray HR=1.20, 95% CI: 0.87-1.67; Tothill HR=0.87, 95% CI: 0.45-1.66) (Figure S2).

Tumor debulking surgery is one of the principal treatments for HGSOC. Surgery outcomes are traditionally classified as optimal (only residual disease is <1cm in diameter) or suboptimal (residual disease 1cm or larger), and optimal debulking is associated with a better prognosis [34]. TME content could affect surgical debulking by making tumors more difficult to remove. Because of this, we wanted to assess if there was a relationship between tumor composition and debulking status. Across all datasets, tumors that had been suboptimally debulked had a higher median fraction of fibroblasts than tumors that had been optimally debulked (Fig. 4E). In contrast, we observed no difference in immune cell fraction between optimally and suboptimally debulked tumors (Fig. 4F).

### Expression programs in the cancer fraction correspond to TME

BayesPrism provides a feature to extract different gene expression programs (“embeddings”) from the estimated malignant fraction using nonnegative matrix factorization (NMF). We performed NMF consensus clustering to select an appropriate number of embeddings (Fig. S3). Based on these results and to match our understanding of the number of subtypes in the stromal data, we ran NMF on each dataset with *k* = 3 (Fig. 5A-C). In all datasets, we found one program expressed highly in samples mapped to cluster 2 (proliferative subtype in *k* = 4). We also found an expression program connected with samples in cluster 3 (immunoreactive and differentiated subtypes in *k* = 4), though less strongly enriched. We did not find any gene expression programs significantly correlated with cluster 1 (mesenchymal subtype at *k* = 4).

**Fig 5.**
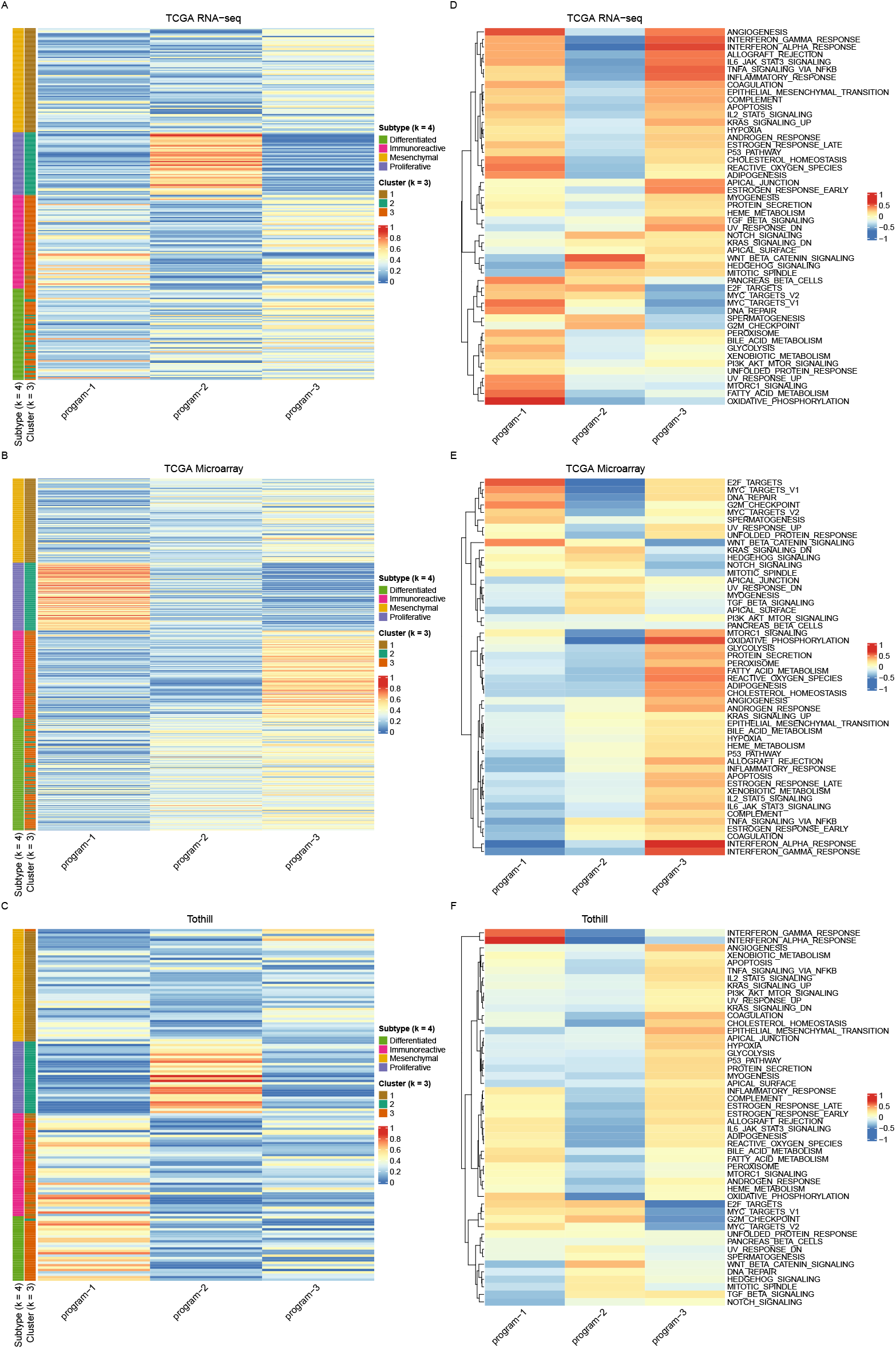
Cancer fraction embeddings across datasets. **(A-C)** Inferred weights of gene expression programs identified using BayesPrism embedding learning at *k* = 3. Samples are ordered according to TCGA subtype. **(D-F)** Gene Set Variation Analysis results comparing the enrichment of MSigDb Hallmark gene sets across tumor embeddings.

We next checked the cancer-specific expression programs against potentially informative gene signatures. Owing to the lack of high-confidence gene sets associated with the HGSOC subtypes, we opted to compare the expression programs with the Hallmark gene sets from MSigDB [35] (Fig. 5D-F). We found that the gene expression program related to the proliferative subtype tended to have the highest enrichment of gene sets relevant to the cell cycle (G2M checkpoint, mitotic spindle, etc.). The expression program related to the immunoreactive and differentiated subtype tended to have the highest enrichment of interferon response pathways and other cytokine signals.

## Discussion

Deconvolution allows us to systematically evaluate compositional differences in the tumor microenvironment. Identifying how differences in tumor composition affect clinical outcomes like survival requires data from hundreds or even thousands of patients, which is currently not feasible with only single-cell profiling. BayesPrism and other single-cell informed deconvolution methods allow the knowledge gained from a smaller number of single-cell sequenced samples to be applied to existing population-scale resources with bulk transcriptomic profiling.

Here, we demonstrate that using BayesPrism on TCGA’s one-color microarray dataset returns comparable results to its RNA-seq dataset, which will allow for the application of TME analysis to additional bulk transcriptomic datasets. In comparing tumor composition estimates to the TCGA transcriptomic subtypes, we confirm that existing subtype definitions reflect differences in abundance of certain cell types. Most notably, the mesenchymal subtype is associated with a high proportion of fibroblasts. Given our results, we propose a three-subtypes model defined by fibroblast and immune content: 1) fibrotic tumors, which have high fibroblast and also high immune cell content, 2) non-fibrotic immune-high tumors, and 3) non-fibrotic immune-low tumors. We believe this more accurately reflects the biological differences in how HGSOC tumors grow in relationship to their microenvironment. While precise proportional thresholds for defining high fibroblast or high immune content will vary based on a study’s population and sampling criteria, the relative comparisons appear to be preserved across studies.

We observed that tumors with a high proportion of fibroblasts tend to have a slightly worse prognosis compared to less fibrotic tumors. There are a number of possible explanations for the relationship between fibroblast content and survival: for one, the dense, rigid extracellular matrix found in a fibrotic tumor may impede infiltration of the tumor by cytotoxic immune cells or chemotherapeutic drugs [36, 37]. Also, a fibrotic tumor may be more difficult to resect effectively. Here, we offer support for this theory by showing that tumors with suboptimal surgical outcomes tend to have more fibroblasts than those with optimal surgical outcomes.

While immune cell content appears to be integral for defining the TCGA subtypes, we did not identify a relationship between total proportion of immune cells and survival. This is not surprising, given that the TCGA subtypes with the best and worst prognosis (immunoreactive and mesenchymal, respectively [11]) both have a high immune signature. Additionally, most immune cell types can have both immune-promoting or immunosuppressive functions, as in M1/M2 polarization of macrophages [38]. The level of granularity in our deconvolution analysis does not allow us to stratify immune cells to this level; further experiments will be needed to fully characterize how differences in the immune compartment may affect survival. One possible future experiment would be to perform single-cell transcriptomic profiling with CITE-seq [39]; proteomic analysis of cell surface markers could allow for more fine-scale annotation of immune cell types and states and then used as an improved deconvolution reference profile. Further work could also employ spatial transcriptomics platforms to measure immune infiltration within the TME in a spatially-aware manner [40].

Through the BayesPrism embedding learning module, we find evidence that cancer cells express different transcriptional programs based on their microenvironment.

Unsurprisingly, cancer cells from tumors assigned to high-epithelial cluster 1 subtype more readily express gene signatures associated with proliferation. Conversely, cancer cells from tumors with a high immune fraction have more expression of cytokine response pathways, potentially indicating differential responses in cell-cell communication. These results highlight the importance of considering TME heterogeneity in HGSOC at all levels of analysis and eventually in clinical decision-making. We believe that stratifying patients by TME composition may allow for identification of specific tumor vulnerabilities based on cancer cells’ crosstalk with their environment, laying the foundation for novel therapeutics and unlocking the potential of precision medicine for HGSOC.

## Materials and methods

### Datasets

#### Single-cell data

Our primary reference profile for deconvolution was comprised of single-cell RNA-sequencing data originating from 15 HGSOC tumors that were collected at two sites, the University of Pennsylvania (Penn) and the University of Utah (Utah). The Penn samples (n = 8) were collected from patients with HGSOC by the University of Pennsylvania Ovarian Cancer Research Center’s Tumor BioTrust Collection (RRID: SCR 022387). All patients underwent primary debulking surgery and had not received neoadjuvant chemotherapy. Protocol details can be found in [26]. Briefly, the Penn samples were single-cell sequenced in two ways, individually and multiplexed into 2 pools of 4 samples each [26]. One sample, 2428, was excluded due to low total cell counts, but cells assigned to that sample in Pool B were used. The Utah samples (n = 7) were collected and sequenced at Huntsman Cancer Institute. Protocol details can be found in [29]. All samples were sequenced individually.

We also used data from Vázquez-García et al [31] (Gene Expression Omnibus (GEO) ID GSE180661) as an independent reference profile. This dataset was too large to feasibly analyze with BayesPrism, which is an accurate but computationally expensive deconvolution algorithm, in its entirety. We took a random subset of approximately 10% of the total annotated cells in the dataset, 93,204 cells in total (Table 3).

**Table 3.**
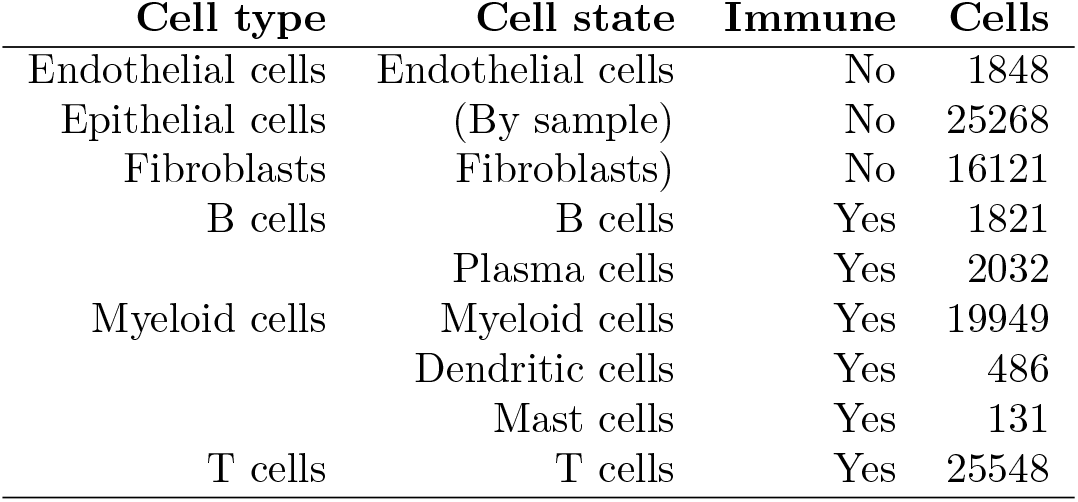
Reference profile data sampled from Vazquez-Garcia et al [31] stratified by cell type and state.

#### Bulk data

We performed deconvolution on three bulk transcriptomic datasets: TCGA RNA-seq [7], TCGA Microarray [7], and Tothill [10], all sourced from the curatedOvarianData R package [41]. See Table 1 for the number of samples per dataset. Both microarray datasets used Affymetrix Human Genome arrays (Affymetrix Human Genome U133 Plus 2.0 Array for TCGA and Affymetrix HT Human Genome U133A Array for Tothill). Crucially, these are one-color arrays, so the one-dimensional (*log*_2_) intensity values can be used as an analog to read counts in RNA-seq data. For these datasets, we used the inverse logarithm values (2^*n*^) as inputs for BayesPrism, which was designed to accept data as counts.

### Data analysis

#### Single-cell processing

All single-cell samples in the HGSOC-Penn/Utah dataset were quantified using Cell Ranger version 6.1.2 and mapped to a GRCh38 reference genome from 10x Genomics (2020-A). We used miQC to filter dead and compromised cells from our single-cell data [42]. We then assigned cell type labels to each sample using a combination of unsupervised clustering and CellTypist [43] as in [26]. Briefly, for each sample, we ran unsupervised clustering and annotated clusters based on marker genes and ran CellTypist in parallel. Cells with concordant assignments based on unsupervised clustering and CellTypist were included in the reference profile for deconvolution. Cells from the multiplexed samples were only included if they were able to be assigned to a cell type and assigned to a donor using the genetic demultiplexing method vireo [44]. Most cells in each sample were able to be assigned as described (Table 4), resulting in a reference profile comprising 79,926 cells across 13 cell types (Table 2).

**Table 4.** HGSOC-Penn/Utah single-cell data stratified by sample.

| Name | Location | Sample Type | Cells |
| --- | --- | --- | --- |
| 2251 | University of Pennsylvania | Individual | 5333 |
| 2267 | University of Pennsylvania | Individual | 5173 |
| 2283 | University of Pennsylvania | Individual | 6432 |
| 2293 | University of Pennsylvania | Individual | 10136 |
| 2380 | University of Pennsylvania | Individual | 5827 |
| 2467 | University of Pennsylvania | Individual | 6397 |
| 2497 | University of Pennsylvania | Individual | 7697 |
| A | University of Pennsylvania | Pooled | 6171 |
| B | University of Pennsylvania | Pooled | 8629 |
| 16030X2 | Huntsman Cancer Institute | Individual | 4606 |
| 16030X3 | Huntsman Cancer Institute | Individual | 933 |
| 16030X4 | Huntsman Cancer Institute | Individual | 3416 |
| 18389X2 | Huntsman Cancer Institute | Individual | 1795 |
| 19459X1 | Huntsman Cancer Institute | Individual | 2632 |
| 19595X1 | Huntsman Cancer Institute | Individual | 618 |
| 19833X1 | Huntsman Cancer Institute | Individual | 2027 |
| 19833X2 | Huntsman Cancer Institute | Individual | 2104 |

BayesPrism requires two levels of cell annotation, that of “cell type” and “cell state” [25]. For most cell types in the HGSOC-Penn/Utah data, we considered these to be synonymous and labeled them identically. However, we used cell state as a label for sample of origin for all malignant/epithelial cells, as recommended by the BayesPrism developers to help capture the known high inter-patient heterogeneity in malignant cells [25, 28]. We also used cell state to better represent the wide phenotypic heterogeneity in T lymphocytes; in this case, “T cell” was treated as the cell type, and cell state was divided into three categories: cytotoxic T cells, helper T cells, and regulatory T cells. BayesPrism only uses cell state information internally and reports all proportion estimates at the cell type level.

For the Vázquez-García data, we used the cell type labels provided by the authors on GEO. We used their annotation hierarchy to define cell type and cell state, with dendritic cells and mast cells considered as subtypes of myeloid cells and plasma cells considered a subtype of B cells (Table 3). Cell state for all malignant cells was defined by sample of origin.

#### Deconvolution

To prepare for deconvolution, we filtered each bulk transcriptomic dataset to genes also found in our single-cell data and excluded all ribosomal and mitochondrial genes. To optimize runtime, we also filtered to include only protein-coding genes. The exact number of genes used for deconvolution of each dataset can be found in Table 1. We built a snakemake [45] workflow to analyze each combination of bulk and single-cell datasets using BayesPrism.

#### Subtypes

Subtyping and cluster assignment for all bulk samples was performed as in [8]. Briefly, we identified the 1,500 genes with the highest Median Absolute Deviation (MAD) in each of five HGSOC microarray datasets: TCGA [7], Mayo [8] (GSE74357), Tothill [10] (GSE9891), Bonome (GSE26712) [46], and Yoshihara [12] (GSE32062). We note that not all datasets that were used to identify MAD genes were included in our deconvolution analyses. Yoshihara and Mayo were sequenced on two-color microarrays, the values of which are not able to be neatly approximated to read counts in order to run BayesPrism, and Bonome was excluded from further analyses due to a lack of survival data. All datasets were obtained from curatedOvarianData [41]. We then ran k-means clustering on the gene expression matrix of the MAD genes for each dataset separately, with *k* = 2, 3, and 4. Consensus clusters were derived from 100 independent clustering runs. We used significance analysis of microarray (SAM) analysis [47, 48] to achieve consistent cluster numbering across datasets and to map to the original TCGA nomenclature (immunoreactive, proliferative, mesenchymal, differentiated) at *k* = 4.

#### Clinical outcomes

We obtained survival and other clinical data for all bulk transcriptomic datasets from the curatedOvarianData R package [41]. Overall survival was measured in months. Age at initial diagnosis was binned into groups of 5 years between 40 and 75 (i.e. 0-39, 40-44, 45-49, …, 75-110). Tumor stage was grouped into stage I, stage II/III, and stage IV. Debulking status was assigned as optimal (only residual disease is *<*1cm in diameter) or suboptimal (residual disease 1cm or larger). Debulking status was missing for approximately 10% of cases across datasets (29 in TCGA RNA-seq, 44 in TCGA microarray, 17 in Tothill) and these samples were excluded from analysis. We visualized overall survival across datasets using Kaplan-Meier curves and generated Cox proportional hazards (PH) models to calculate hazard ratios (HR) using the R package survival [49, 50].

#### Cancer expression programs

For each bulk dataset, we ran NMF on the estimated cancer fraction expression returned by BayesPrism. We used the R package NMF [51] to calculate factorization rank metrics for a range of possible *k* values from 2 to 12 and generate a consensus map for each value of *k*. Based on these results, we decided to run NMF with *k* = 3 and *k* =4. We ran NMF on each bulk dataset using BayesPrism’s *run*.*embedding*.*nmf* function for 40 cycles. We plotted the embedding weights (the relative expression of each embedding gene signature) for each sample and compared them to the subtype clusters. We performed Gene Set Variation Analysis [52] to assess how the embedding gene signatures aligned with the MSigDB Hallmark gene pathways [35].

## Availability of data and materials

The HGSOC-Penn/Utah single-cell dataset is available in GEO (processed gene count tables) under accession numbers GSE158937 and GSE217517, and in the Database of Genotypes and Phenotypes (dbGaP) (raw FASTQ files) under accession phs002262.v3.p3. The Vázquez-Garía single-cell dataset is available in GEO under accession number GSE180661. The code for all the analyses performed in this paper is available at https://github.com/greenelab/hgsoc_deconvolution under a BSD-3-Clause license.

## Acknowledgments

The authors would like to thank Ronny Drapkin, Dalia Omran, and other members of the Penn Ovarian Cancer Research Consortium for contributing and preparing samples for use in the HGSOC-Penn/Utah dataset. Thanks also to Jake Crawford for code review.

## Author Contributions

- AAH: Conceptualization, methodology, formal analysis, investigation, data curation, software, visualization, writing – original draft
- NRD: Data curation, formal analysis, software, writing – review & editing
- MEB: Data curation, writing – review & editing
- LMW: Data curation, writing – review & editing
- JG: Project administration, resources, writing – review & editing
- JAD: Conceptualization, funding acquisition, writing – review & editing
- SCH: Conceptualization, funding acquisition, writing – review & editing
- CSG: Conceptualization, funding acquisition, writing – review & editing

## Funding

AAH, LMW, JAD, CSG, and SCH were supported by the National Cancer Institute (NCI) from the National Institutes of Health (NIH) grant R01CA237170. MEB received support from the National Cancer Institute (K00 CA212222) and the Karin Grunebaum Cancer Research Foundation. Generation of the data used in this study was also funded by a Hopkins-Penn Ovarian Cancer SPORE Developmental Research Program award, P50 CA228991. The funders had no role in study design, data collection and analysis, decision to publish, or preparation of the manuscript.

## Competing Interest Statement

The authors declare that they have no competing interests.

## Supplementary Figures

**Fig S1.**
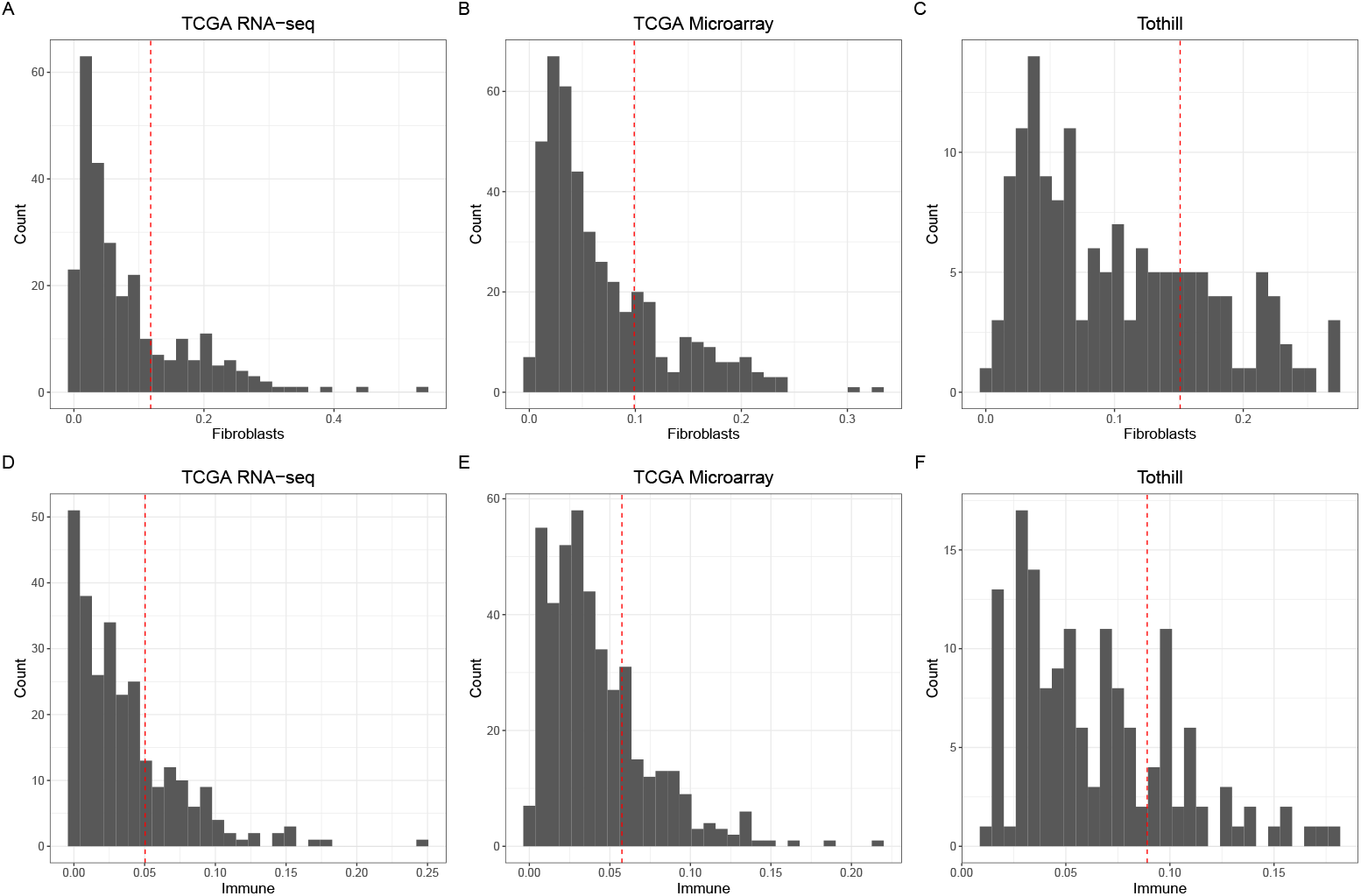
Tumor composition and clinical outcomes. **(A-C)** A histogram of the proportion of fibroblasts in each bulk transcriptomic dataset. The red dashed line indicates where the third quartile is; all samples above are considered “high fibroblast” for survival analyses. **(D-F)** The proportion of immune cells in each datasest, with a red dashed line at the third quartile.

**Fig S2.**
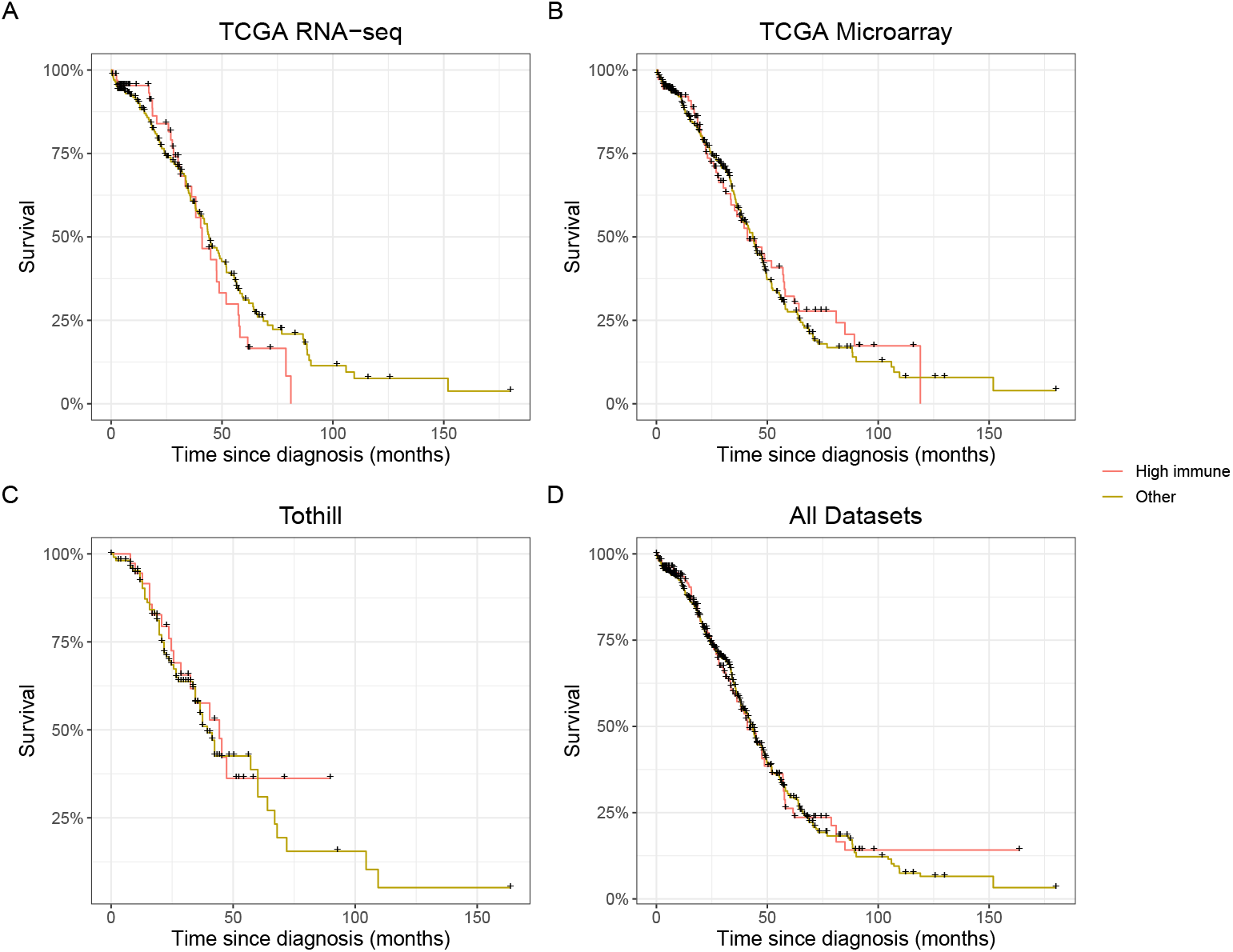
Immune composition and survival. **(A-D)** Kaplan-Meier survival curves stratified between high immune tumors (top quartile of immune content per dataset) and non-high immune (all other quartiles).

**Fig S3.**
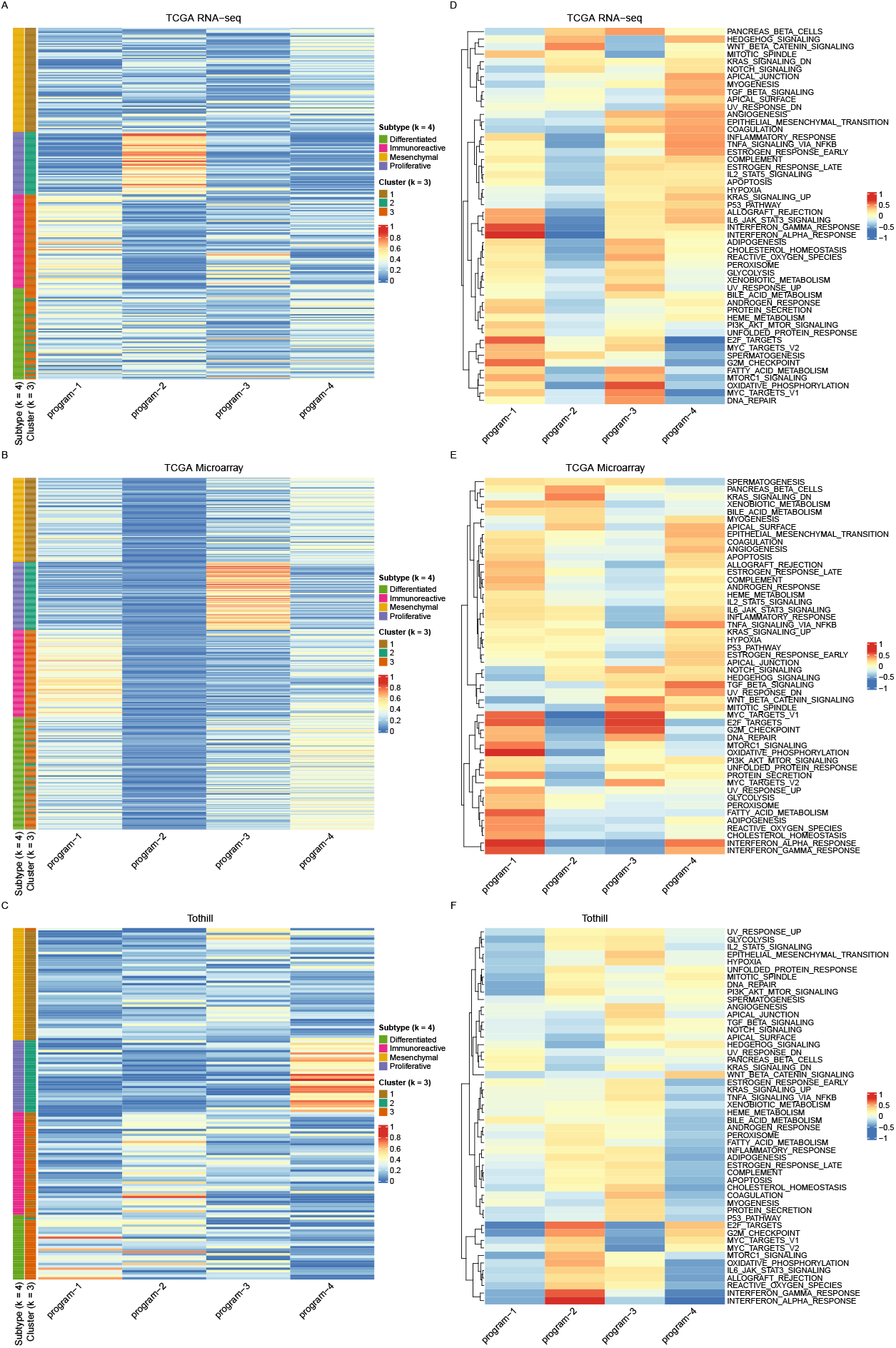
Cancer fraction NMF consensus clustering results. **(A-C)** A map of consensus clustering from *k* = 2 to 6 and cophenetic correlation results for *k* = 2 to 12 for the TCGA RNA-seq **(A)**, TCGA Microarray **(B)**, and Tothill **(C)** datasets.

## Notes

https://github.com/greenelab/hgsoc_deconvolution

### Competing Interest Statement

The authors have declared no competing interest.

